# A novel AAA+ ATPase required for sporulation and stress response in *Bacillus anthracis*

**DOI:** 10.1101/2025.06.16.659970

**Authors:** Nitika Sangwan, Ankur Bothra, Andrei P. Pomerantsev, Aakriti Gangwal, Rasem Fattah, Mahtab Moayeri, Qian Ma, Sundar Ganesan, Chetkar Chandra Keshavam, Renu Baweja, Uma Dhawan, Stephen H. Leppla, Yogendra Singh

**Affiliations:** Department of Biomedical Science, Bhaskaracharya College of Applied Sciences, University of Delhi; Microbial Pathogenesis Section, Laboratory of Parasitic Diseases, National Institute of Allergy and Infectious Diseases, Bethesda, MD, USA; Department of Zoology, University of Delhi, Delhi, India; Biological Imaging Section, Research Technologies Branch, National Institutes of Allergy and Infectious Diseases, National Institutes of Health, Bethesda, MD, USA; Department of Biochemistry, Shivaji College, University of Delhi; Delhi School of Public Health, Institution of Eminence, University of Delhi

**Author notes:** Authors contributed equally (Nitika Sangwan, Ankur Bothra). Corresponding Authors: Yogendra Singh, Advisor, Delhi School of Public Health, Institution of Eminence, University of Delhi, India; Email ID; Stephen H. Leppla, Chief, Microbial Pathogenesis Section, Laboratory of Parasitic Diseases, NIAID, NIH, Bethesda, USA; Email ID; Uma Dhawan, Professor, Department of Biomedical Science, Bhaskaracharya College of Applied Sciences, University of Delhi, India; Email ID.

**Keywords:** AAA+ ATPase, BAS PrkA, *Bacillus anthracis*, Sporulation, Osmotic stress, Anthrax

## Abstract

AAA+ proteins function as molecular machines that utilize ATP to perform diverse cellular functions, including protein homeostasis, stress regulation, and cell cycle/developmental processes. In this study, we identified a novel AAA+ ATPase BAS PrkA in *B. anthracis* Sterne 34F2 which has 88 % protein homology to *Bacillus subtilis* PrkA. Conserved domain analysis confirms BAS PrkA has an N-terminal AAA+ ATPase domain with characteristic Walker A and Walker B motifs and a conserved secondary region of homology (SRH) domain, along with a C-terminal cAMP-dependent protein kinase domain. Based on Alpha Fold3 predicted structure, we classified BAS PrkA as part of Clade III of the AAA+ superfamily. Contrary to the reported enzymatic activity in *B. subtilis* PrkA, we observed that BAS PrkA has negligible protease and kinase activity under *in-vitro* conditions. Nonetheless, BAS PrkA plays a significant role in regulating sporulation. It is temporally expressed during Stages II to VI during sporulation. A null mutant of BAS PrkA exhibits severe sporulation defects, with reduced spore viability, and down regulation of genes related to spore-coat formation. These phenotypes were restored in a complementation strain expressing BAS PrkA ectopically. Additionally, the null mutant strain showed compromised growth under ionic-osmotic stress conditions. Analysis of the BAS PrkA interactome revealed enrichment of two proteins, ProA and EzrA, that are implicated in osmotic stress response and the sporulation process, respectively. These findings show that BAS PrkA plays a critical role in sporulation and osmotic stress response in *B. anthracis*.

## Introduction

The life cycle of *Bacillus* species consists of three phases: vegetative growth, sporulation, and germination. During the vegetative phase, the bacteria grow and divide actively [1]. When growth conditions become unfavorable, particularly due to nutrient limitation or environmental stress, the bacteria initiate sporulation, forming highly resistant endospores. These endospores are metabolically dormant, allowing the bacteria to survive under adverse conditions. Once conditions are favorable, the endospores undergo germination and re-enter the vegetative phase to resume growth and division.

Protein quality control and homeostasis are essential during these transitions between vegetative bacteria and spores, with ATP-dependent AAA+ (ATPases Associated with diverse cellular Activities) proteins playing key roles [2, 3]. In *Bacillus subtilis* and *Bacillus anthracis*, members of the Clp, Lon, and FtsH families represent well-characterized AAA+ proteins involved in various cellular processes including stress responses and sporulation [4–10]. Given the key roles undertaken by AAA+ proteins in *Bacillus spp.*, studying novel AAA+ domain-containing proteins could reveal new insights into their physiological roles in *B. anthracis*.

Here, we show that *B. anthracis* BAS0518 (BAS PrkA, now onwards) is a homolog of *B. subtilis* PrkA with a predicted AAA+ ATPase domain that exhibits negligible protease activity *in vitro*. This contrasts with *B. subtilis* PrkA (BS PrkA) which was reported to have ATP-dependent protease activity *in-vitro,* although the activity was quite limited [11]. Using a BAS PrkA deletion mutant, we demonstrate that BAS PrkA is not required for normal growth during the vegetative phase. However, under osmotic stress conditions, the survival of the BAS PrkA deletion mutant is severely compromised. We further reveal that BAS PrkA is not produced in actively growing vegetative bacteria and appears only during Stage-II of sporulation, and deletion of BAS PrkA leads to reduced spore viability. Additionally, BAS PrkA deletion results in down regulation of several sporulation-related genes at RNA level, including *cotE* and *sigK* in the log phase, *sigK* and *spoIID* in the late exponential/early stationary phase and *gerE* in the late stationary phase of bacterial growth.

We identified that loss of BAS PrkA significantly alters the spore-coat layer, resulting in low-viability and heat-sensitive spores. Lastly, through PrkA interactome identification, we identify potential BAS0518 interacting partners, ProA and EzrA, alteration of which may contribute to the observed phenotypes related to stress response and sporulation, respectively. Together, these findings suggest that PrkA is required for efficient sporulation and stress response.

## Results

BAS PrkA, a sequence ortholog of *B. subtilis* PrkA, has no detectable enzymatic activity BAS0518 (BAS PrkA) from *B. anthracis* (locus WP_268553006.1) is annotated as a serine/threonine protein kinase (PrkA family) based on sequence similarity to *B. subtilis* PrkA, a putative kinase previously identified through its distant homology to eukaryotic kinases and in vitro phosphorylation of a 60-kDa protein [14]. Detailed structural domain analysis of BAS PrkA protein revealed the presence of two domains, an N-terminal domain related to the AAA+ (ATPases associated with diverse cellular activities) superfamily, and a C-terminal domain having distant homology to (cAMP)-dependent protein kinases (Figure 1A).

**Figure 1:**
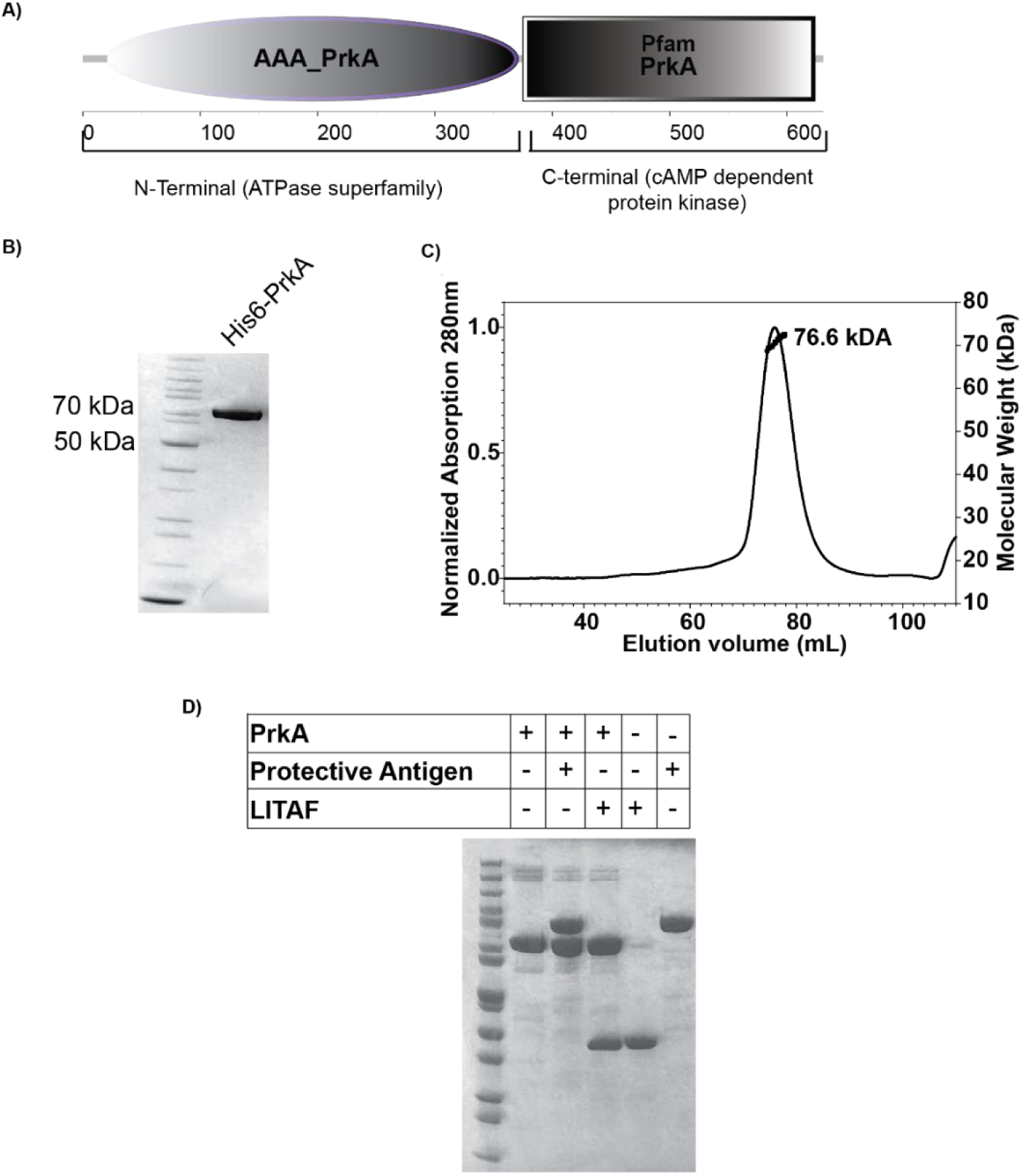
PrkA has no detectable protease activity. **(A)** Domain analysis of BAS PrkA reveals that the N-terminal region of the BAS PrkA protein belongs to the AAA+ superfamily of ATPases, while the C-terminal region is homologous to cAMP-dependent kinases. **(B)** Coomassie stained SDS-PAGE showing the purified His6 PrkA protein. **(C)** Confirmation of purified His6 PrkA protein by size-exclusion chromatography coupled with multi-angle light scattering (SEC-MALS). The elution profile shows His6 PrkA at a molecular mass of 76.6 kDa, using buffer containing 5 mM CHAPS and 1 mM TCEP. **(D)** Coomassie-stained SDS-PAGE for measuring (non)proteolysis of LITAF and PA by monomeric His6 PrkA at 24 hours, incubated at 37°C. His6 PrkA and substrates (LITAF, PA) were incubated separately at 37°C as controls. Enzyme and substrates were used at amount of 2 µg per lane.

Supplementary figure-1A describe the detailed domain analysis of BAS PrkA. In the N-terminal part of the protein sequence, we found the conserved Walker A and Walker B motifs, which are characteristic of ATPases known for ATP binding and hydrolysis [12]. Other conserved motifs include sensor motifs that interact with gamma-phosphate of bound ATP and thus senses the nucleotide binding domain in substrates. In the C-terminal part of protein, 11 conserved motifs typical of the Hanks-type kinases are present [13]. Interestingly, the P-loop, which is important for ATP binding in Hank-type kinases, is absent in the C-terminal part of the PrkA protein sequence. As a result, His6-PrkA exhibited no detectable autophosphorylation activity, suggesting a lack of intrinsic kinase function. However, His6-PrkA did undergo low-level phosphorylation by catalytic domain of another well-characterized *B. anthracis* Ser/Thr/Tyr kinase (STPK), PrkC (PrkC^cat^-GST) [14]. (Supplementary figure-1B, C).

Based on structure modelling using Alpha Fold 3, we classified BAS PrkA as part of Clade III of the AAA+ superfamily. Clade III of the AAA+ superfamily of proteins is characterized by a pore-loop 1 inserted between β-sheet 2 (β2) and β-sheet 3 (β3) of the core AAA+ domain [15]. This feature was identified in the predicted monomeric structure of BAS PrkA (Supplementary figure 2A-B). Additionally, the predicted hexameric structure suggests the presence of several aromatic residues in pore loop 1 of BAS PrkA, like what was observed in the representative Clade III structure of YME1 (PDB: 6AZ0) [16] (Supplementary figure 2C-E).

A previous study on BS PrkA in *B. subtilis* suggested weak sequence similarity to the Lon-proteases and detected protease activity using α-casein as the substrate [11]. This led us to investigate the protease activity of the *B. anthracis* BA PrkA. To characterize BAS PrkA of *B. anthracis*, we cloned the BAS0518 (BAS PrkA) gene into *Escherichia coli* expression vector pPro-Ex-Htc, which adds a hexa-histidine tag (His6) at the N-terminus of the protein. The protein was over-expressed and purified from *E. coli* (BL21) strain using Ni-NTA affinity chromatography, resulting in a single band of 76.7 kDa on SDS-PAGE (Figure 1B).

A detailed MS/MS analysis on the initial purification resulted in protein with minor (<0.1 %) contamination of an amido-hydrolase (locus tag #ECBD_2279) of *E. coli*. This contamination may have contributed to the weak protease activity we observed using a fluorescent assay with BODIPY FL-Casein as the substrate, the same substrate used in the study of BS PrkA (Supplementary figure-3B) [11]. To circumvent this issue, further purification was performed using buffers containing 5 mM CHAPS along with 1 mM TCEP, which yielded a stable monomeric form of the protein. This was confirmed using size-exclusion chromatography coupled with multiple angle light scattering (SEC-MALS) (Figure 1C). To confirm the identity of the purified protein, we performed intact-mass analysis using mass-spectrometry, which revealed a single peak of 76,639 Da, consistent with the calculated mass of the His6 PrkA protein. This confirmed the efficacy of our protein purification, and this protein was used for further biochemical characterization.

To assess the protease activity of His6 PrkA, we used LITAF and anthrax Protective antigen (PA) as substrates and analysed cleavage on Coomassie-stained SDS-PAGE. The selection of LITAF and PA was based on observations that they contain long, unfolded sequences which make them sensitive substrates for cleavage analysis, as represented in supplementary figure-3A [17, 18]. As shown in supplementary figure-3C, using SDS-PAGE we observed a weak protease activity of purified His6 PrkA (containing the putative amido-hydrolase) on the LITAF and PA proteins. However, the purest monomeric form of His6 PrkA did not cleave these substrates even after 24 hours of incubation (Figure 1D). Based on the AlphaFold3 prediction (Supplementary figure 2C-E), BAS PrkA is structurally similar to the YME1 protease of *Saccharomyces cerevisiae* (PDB-6AZ0) but lacks the active site residues necessary to perform protease activity. These *in-vitro* findings leave the biological role of *B. anthracis* unresolved.

### Physiological role of BA PrkA in *B. anthracis* stress response

Since BS PrkA in *B. subtilis* is suggested to be involved in sporulation, we hypothesized that BAS PrkA may have a similar role in *B. anthracis* [19].To understand the physiological role of BAS PrkA in *B. anthracis* Sterne 34F2 (BAS), we created an unmarked gene deletion mutant using a method described previously [20] (Figure 2A, B). The mutant strain (BAS Δ*prkA*) was assessed under normal and salt stress condition to determine its physiological role.

**Figure 2:**
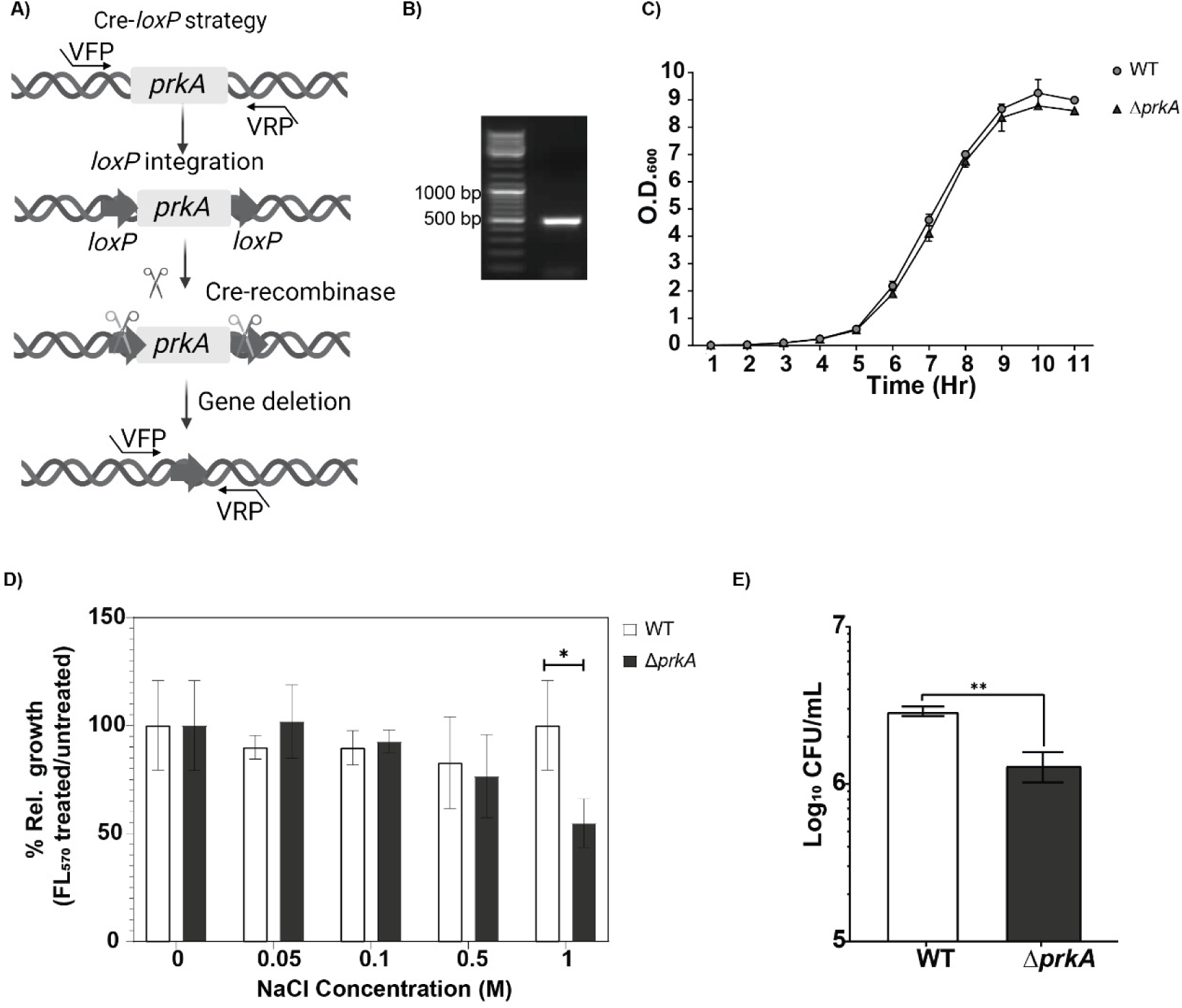
Deletion of BAS *prkA* does not affect growth under normal conditions but causes growth defects under ionic osmotic stress. **(A)** Schematic diagram representing Cre-*loxP* based homologous recombination strategy for BAS *prkA* gene deletion. VFP and VRP are BAS *prkA* gene flanking primers. **(B)** Agarose gel showing PCR confirmation of BAS *prkA* gene deletion. The deletion is verified by the presence of a 495 bp PCR amplicon generated using primers flanking the BAS *prkA* gene (lane 2). Lane 1: 1 kb DNA ladder. **(C)** Growth curves of BAS wild-type (WT) and BAS *ΔprkA* strains in LB medium. Values represent the mean optical density at 600 nm (OD_600_) ± standard deviation (SD) from n = 3 replicates. **(D)** Bar graph representing mean ± SD of the percent relative growth (y-axis) of BAS WT and BAS *ΔprkA* strains in LB medium supplemented with varying concentrations of NaCl. Growth is represented as a percentage of the maximum growth observed in the absence of NaCl (x-axis). n = 2 experiments. **(E)** Bar graph showing mean ± SD of colony-forming units per ml (CFU/ml) of BAS WT and BAS *ΔprkA* strains surviving in LB medium supplemented with 1 M NaCl. N = 2 experiments. Asterisks indicate statistical significance calculated using a two-tailed Student’s t-test (*p < 0.05, **p<0.01).

Under nutrient-rich conditions (Luria-Bertani broth), the BAS Δ*prkA* exhibited a growth profile like the BAS WT (wild-type) strain over ∼10 hours. (Figure 2C). Among many different players involved in stress responses, AAA+ superfamily of proteins has been shown to be critical for different stress coping mechanisms across species [21]. Hence, we examined the role of BAS PrkA under different stress conditions, including ionic-osmotic stress (1 M NaCl), non-ionic osmotic stress (20 % sucrose), and oxidative stress (10 mM H_2_O_2_). Bacterial growth under these stresses was measured using resazurin dye to compare growth of untreated and treated cells. For, osmotic stress conditions, we used different concentrations of NaCl (0.05, 0.1, 0.5 and 1 M) and observed a significant decrease in the fluorescence at 1 M NaCl, whereas no differences were observed under other stress conditions (Supplementary figure 4A-B). To further validate the difference observed in cell viability at 1 M NaCl, we counted colony-forming units per millilitre (CFU/mL) of BAS WT and BAS *ΔprkA* strains in 1 M NaCl and observed about a 2.1-fold difference in CFU between the two strains (Figure 2D-E). A significant reduction in the growth of BAS *ΔprkA* show the role of PrkA in osmotic stress condition.

### BAS PrkA is abundantly expressed during sporulation and is required for spore viability

As shown earlier, PrkA plays a significant role in sporulation in *Bacillus subtilis* [11, 19]. Therefore, we assessed the involvement of BAS PrkA in the sporulation process in *B. anthracis* Sterne 34F2. To test this, we first assessed the expression of BAS PrkA during vegetative growth and at different stages of sporulation (Stages 0 to VII). BAS PrkA expression was absent in vegetative cells but began increasing at Stage II, coinciding with the formation of the polar septum. Its expression gradually decreased as spores matured (Figure 3A), suggesting a role for PrkA in equipping *B. anthracis* to complete sporulation.

**Figure 3:**
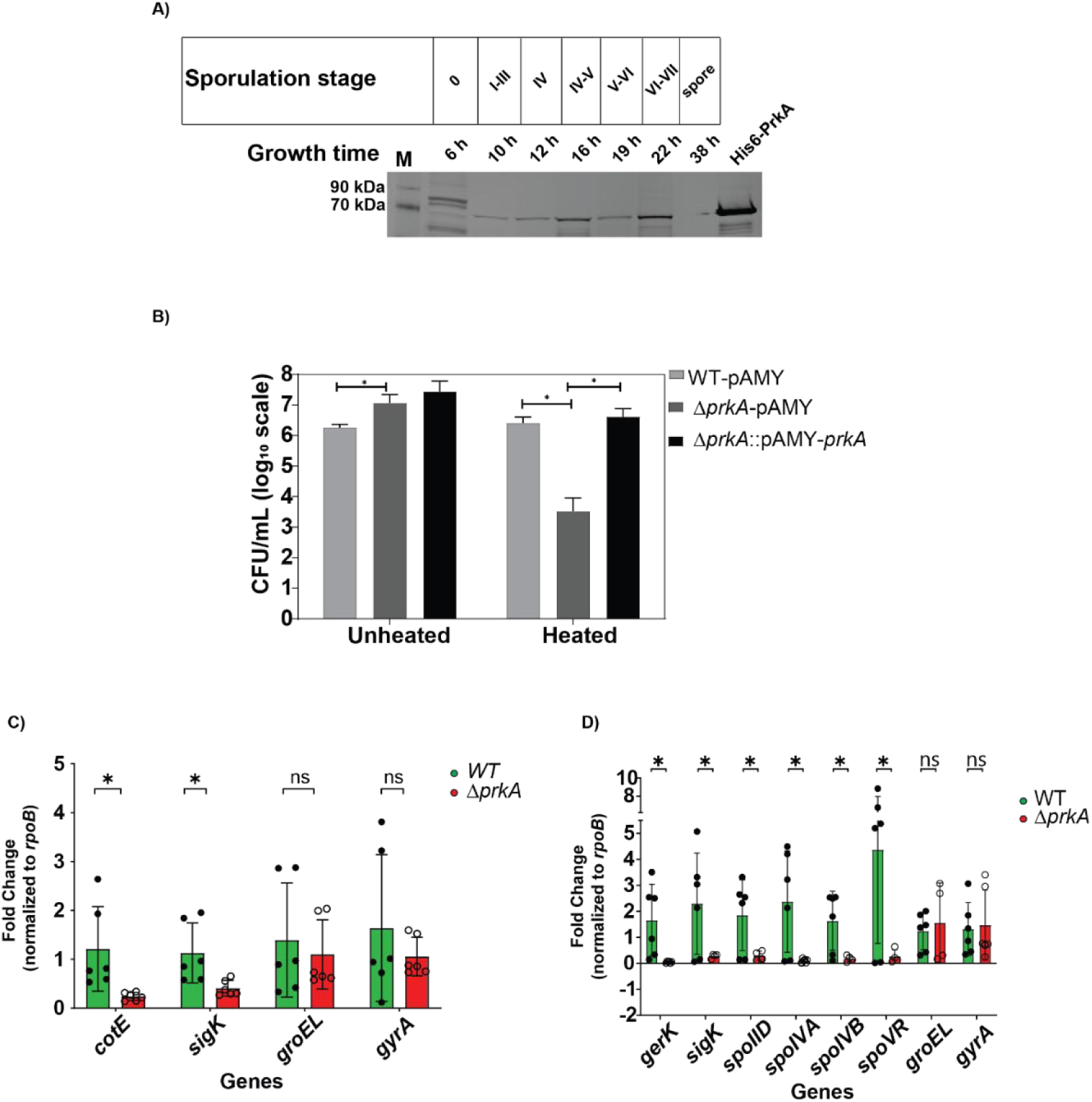
BAS *prkA* is highly expressed during sporulation stages, and its deletion reduces spore viability and alters sporulation gene expression. **(A)** Temporal expression of BAS PrkA protein at various time points corresponding to different stages of sporulation. Growth time and corresponding sporulation stages are mentioned on the top. Lane M: protein ladder, lane 8: purified His6 PrkA protein (positive control). Data are representative of N = 2 experiments. Protein in each lane is normalized to total protein concentration estimated using BCA method. **(B)** Bar graph depicting mean ± SD of sporulation efficiency as colony-forming units per millilitre (CFU/mL) for BAS WT-pAMY, BAS *ΔprkA*-pAMY, and BAS *ΔprkA*::pAMY-*prkA* (complemented) strains. Values represent the mean CFU/mL ± standard deviation (SD) from N = 5 experiments. **(C, D)** Bar graphs showing mean ± SD of relative expression of sporulation genes during early (**C**, Stage 0–III) and late (**D**, Stage V–VI) stages of sporulation. The expression levels were normalized to *rpoB* (housekeeping gene). N = 3 experiments. Asterisks indicate statistical significance calculated using a two-tailed Student’s t-test (*p < 0.05).

Given this stage-specific expression of BAS PrkA during sporulation, we next examined its effect on sporulation efficiency. As a control, a complemented strain of BAS *ΔprkA* was created by ectopically expressing BAS PrkA under an IPTG-inducible promoter (Table 1). Sporulation efficiency was measured by quantifying spore counts in BAS WT-pAMY, BAS *ΔprkA*-pAMY, and BAS*ΔprkA*::pAMY-*prkA* (complemented) strains under both unheated and heated (75°C) conditions. The WT strain produced approximately 6 × 10⁶ CFU/mL of mature spores after five days of nutrient starvation (Figure 3B). However, the BAS *ΔprkA* strain showed a significant 3-log_10_ reduction in mature spores. This sporulation defect was rescued in the complemented strain, confirming a major role of BAS PrkA in sporulation. Similar observations were made in *B. subtilis ΔprkA* mutants, which also showed reduced sporulation efficiency [11].

**Table 1:**
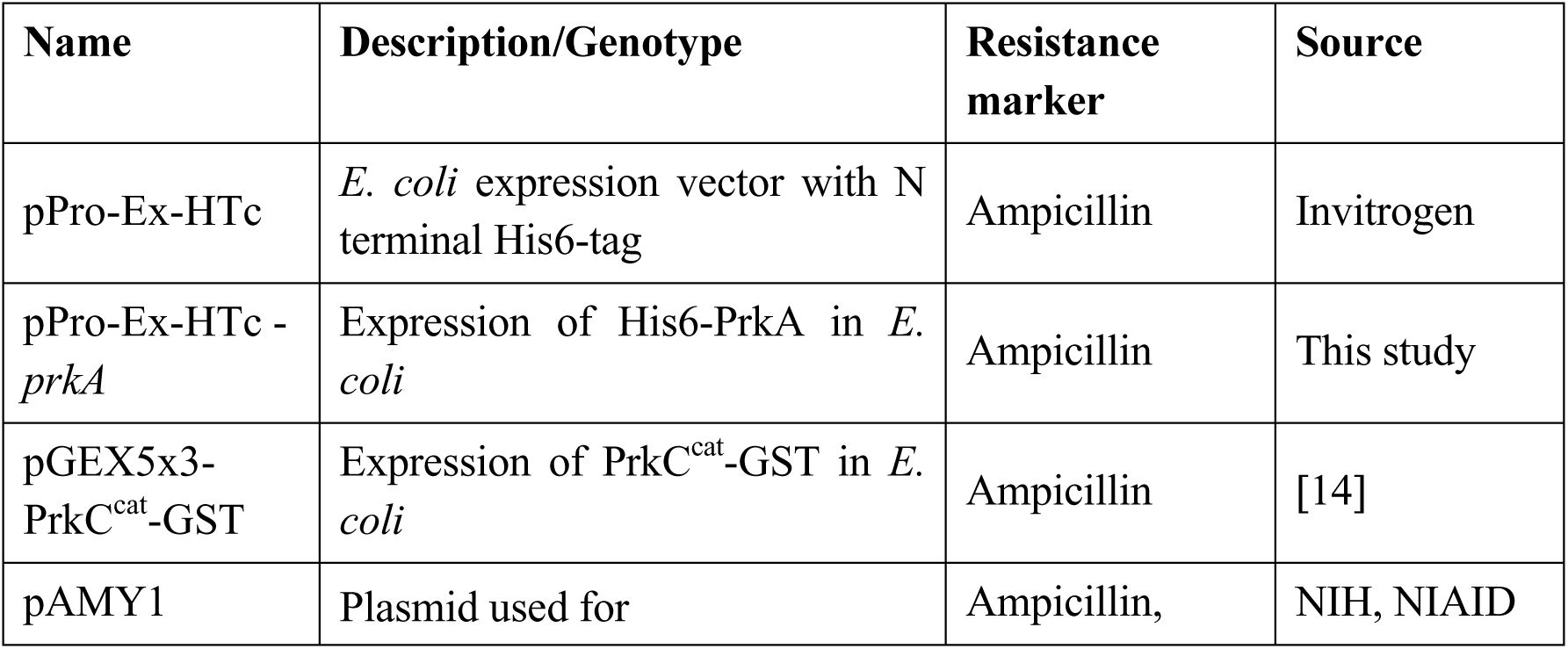
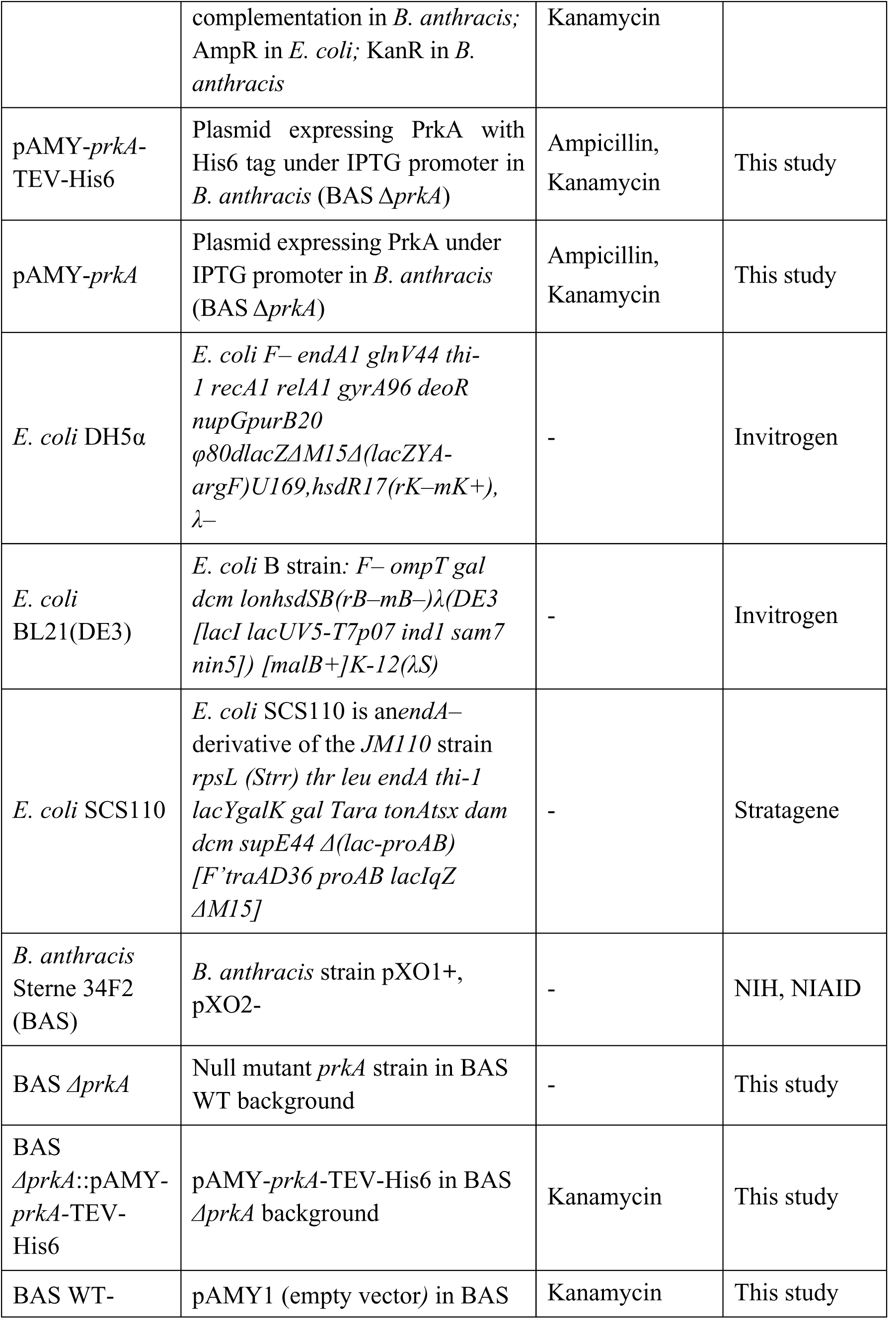
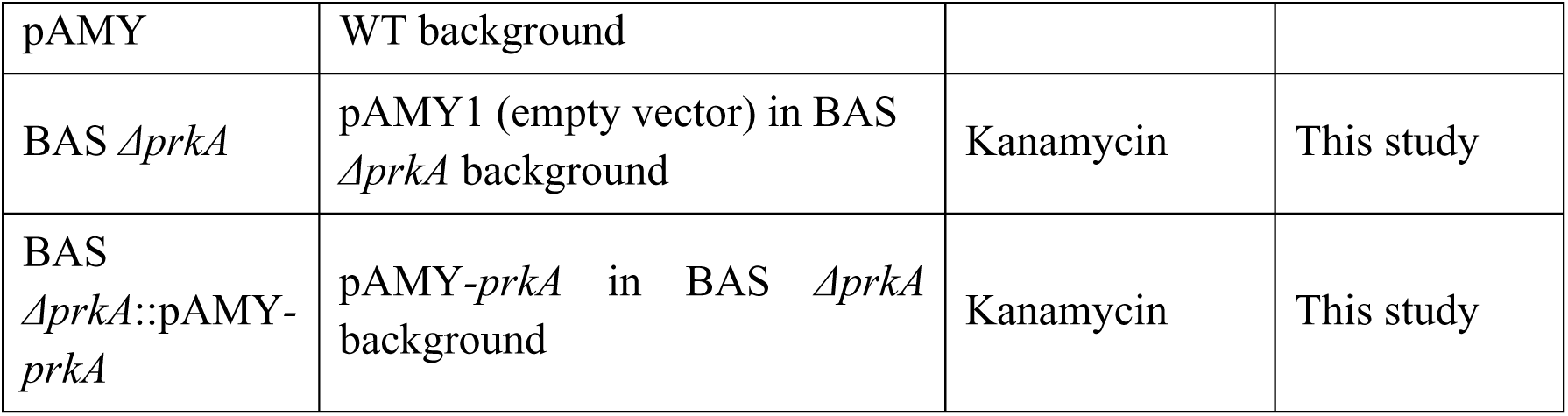
List of plasmids and strains used in this study.

The sporulation pathway is well-characterized in *Bacillus* species, where sigma factors *sigF*, *sigG*, *sigE*, and *sigK* regulate sporulation-related genes in a stage-specific manner [22]. For example, *sigF*, expressed in the forespore, regulates *spoIVB*, while *sigE*, expressed in the mother cell, regulates *spoVB* and *spoIID*. Similarly, *sigG* (forespore) controls *gerD*, and *sigK* (mother cell) regulates *gerE*.

Based on observations of its expression during sporulation and its role in sporulation, we analysed the gene expression of key sporulation genes across early and late stages of sporulation. The expression of genes regulated by several sigma factors (*sigF*, *sigG*, *sigE*, and *sigK*) was evaluated [22]. At the early stages of sporulation (Stages 0-III), we observed significant reductions in the expression of *cotE* (4-fold) and *sigK* (2.5-fold) in the BAS Δ*prkA* strain (Figure 3C, Supplementary figure 5A). At a later stage of sporulation, overlapping with Stage V-VI, other sporulation genes such as *spoIID*, *spoIVA*, *spoIVB*, *spoVR*, and *gerK*, also exhibited reduced expression, suggesting a pleiotropic effect for BAS PrkA in regulating sporulation (Figure 3D, Supplementary figure 5B). In BAS *ΔprkA*, the reduced expression of *sigK* are likely to impact their downstream target genes, disrupting the sporulation pathway, as shown previously [19]. This dysregulation of sporulation genes provides a mechanistic basis for the heat-sensitive phenotype observed in BAS *ΔprkA* spores.

### BAS PrkA is required for mature spore formation

To understand why BAS *ΔprkA* has difference in sporulation, we tracked the progression of sporulation in the mutant strain by staining sporulating bacteria with the cell-permeable (but spore-impermeable) hexidium iodide nucleic acid stain. Previous studies have shown that the bacterial chromosome replicates, condenses, and segregates into the spore and mother cell during sporulation [23]. Therefore, to monitor the differences in sporulation stages, we expect changes in the diffusion pattern and position of the condensed DNA after staining with hexidium iodide at different stages of sporulation. Fluorescence microscopy revealed that BAS *ΔprkA* cells progressed through the same sporulation stages as the WT strain (Stages I to VI) and formed spores (Figure 4A). However, BAS *prkA* produced 3-log lower number of mature spores (Figure 3B), suggesting a role for PrkA in sporulation.

**Figure 4:**
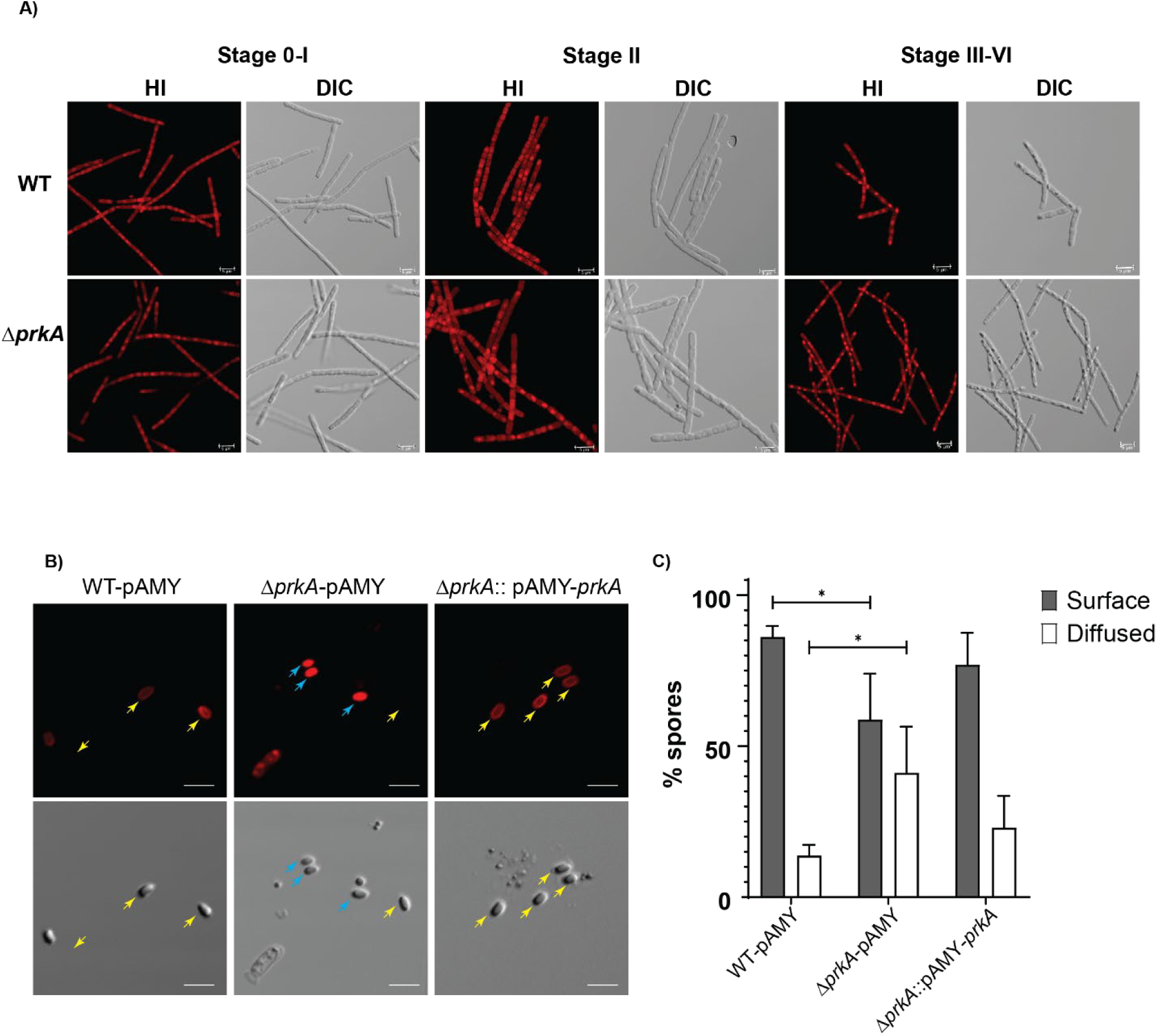
Deletion of BAS *prkA* affects mature spore formation: **(A)** Representative hexidium iodide [HI] stained and differential interference contrast [DIC] images of BAS WT and BAS *ΔprkA* at different sporulation stages (6 hours to 19 hours; Stage 0 to Stage VI); Scale bar (2µm) is depicted in all images. N = 3 biological replicates. **(B)** Representative HI stained and DIC images of BAS WT, BAS *ΔprkA*, and BAS *ΔprkA*::pAMY-*prkA* spores. Scale bar = 2 µm. Yellow arrows indicate spores with non-permeabilized surface staining (hexidium iodide), and blue arrows indicate spores with diffused, permeabilized hexidium iodide staining. Data are from N = 3 biological replicates. **(C)** Bar graph showing mean ± SD of the percentage of spores exhibiting diffused staining versus surface-localized staining. Data are from >2000 spores counted across three biological replicates. Asterisks indicate statistical significance calculated using a two-tailed Student’s t-test (*p < 0.05).

To identify the reason for heat sensitivity in BAS *ΔprkA* spores, we investigated whether the loss of PrkA disrupted spore-coat formation, given the significant reduction in the expression of spore-coat genes (*cotE*, *spoIVA*, and *spoIVB*). The integrity of the spore-coat was tested using spore-impermeable hexidium iodide on the BAS *ΔprkA* spores. As shown in Figures 4B-C, the spores from the BAS *ΔprkA* strain showed significant diffusion of hexidium iodide through their spore-coat. This phenotype was not present in the WT and the complemented (BAS *ΔprkA*::pAMY-*prkA*) strains, suggesting a weak spore-coat layer formation in the BAS *ΔprkA* strain. A weak spore coat results in immature spore formation with a heat-sensitive phenotype, indicating a defect in the structural formation of spores.

### BAS PrkA interactome consists of proteins involved in septum formation, stress response and metabolic genes

To uncover the mechanism by which BAS PrkA influences sporulation in *B. anthracis* Sterne 34F2, we identified its intracellular interactors. Using co-affinity purification followed by mass-spectrometric analysis of whole-cell lysates from a *B. anthracis* Sterne 34F2 strain constitutively expressing His6-PrkA, we identified several BAS PrkA-associated proteins (experimental flow-chart in Figure 5A). A total of 22 co-purifying proteins were identified as potential direct or indirect interactors of PrkA, with a comprehensive list provided in the supplementary table. These interactors were categorized into five major functional groups: cell division/septation, metabolic pathways, stress response, gene regulation, and transporters (Figure 5B).

**Figure 5:**
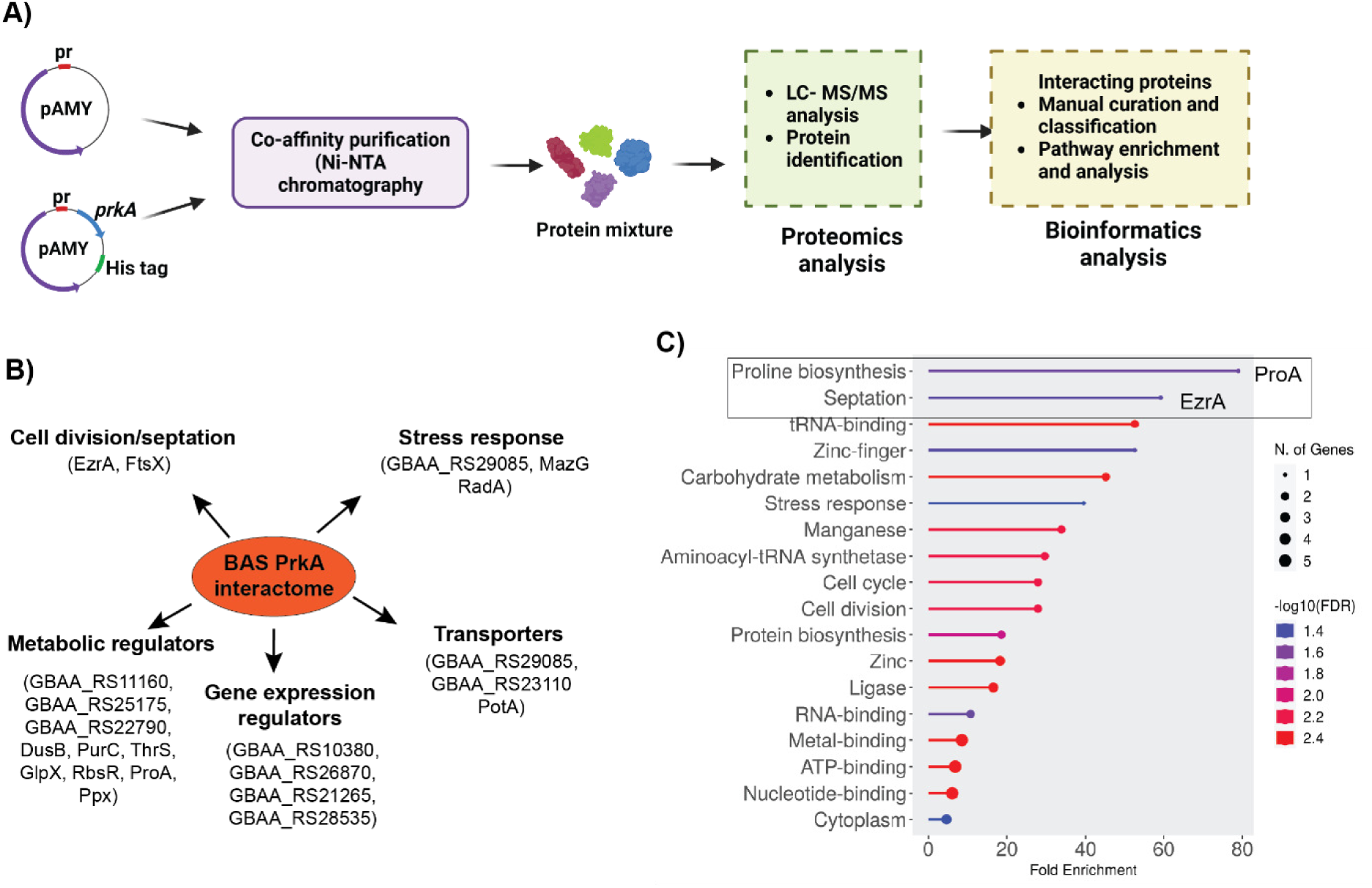
BAS PrkA interactome analysis reveals binding partners potentially linked to compromised spore viability and osmotic stress sensitivity. **(A)** Schematic representation of the co-affinity purification strategy used to isolate the BAS PrkA interactome, followed by mass spectrometry analysis and subsequent interactome characterization. **(B)** Broad classification of the 22 proteins identified as BAS PrkA interacting partners. A detailed table listing these proteins is provided in the supplementary materials. **(C)** Pathway enrichment analysis showing statistically enriched pathways in the BAS PrkA interactome. Notably, EzrA and ProA are highlighted, as they are associated with sporulation and osmotic stress response according to the literature.

Pathway enrichment analysis performed using ShinyGO [24] mapped 11 of the 22 proteins to one or more functional categories (Figure 5C). Among these, ProA and EzrA emerged as the most enriched proteins. ProA, a critical enzyme in proline biosynthesis, has been previously linked to osmotic stress tolerance [25, 26]. EzrA, a key regulator of septation ring formation, plays a crucial role in the sporulation process [27]. Additionally, Ppx, another identified interactor, is implicated in both osmotic stress response and sporulation efficiency [28]. These findings suggest that PrkA exerts its effect on sporulation and stress response by interacting with proteins like ProA, EzrA, and Ppx.

Based on these results, we propose that the observed phenotypes in the BAS0518 *prkA* deletion strain stem from disrupted interactions between BAS PrkA and its interacting partners. Through direct or indirect modulation of these proteins, BAS PrkA may regulate osmotic stress tolerance and the progression of the sporulation process. These findings highlight the central role of BAS PrkA as a regulatory hub connecting stress responses, metabolic adaptation, and cellular processes essential for spore maturation and viability.

## Discussion

In this study, we characterized a novel *B. anthracis* protein, BAS0518 (BAS PrkA), and uncovered its functional role in sporulation and stress adaptation. BAS PrkA contains a putative N-terminal AAA+ ATPase domain and a C-terminal kinase domain, with a strong homology (88 % identical at protein level) to BS PrkA in *B. subtilis*. Our findings highlight that BAS PrkA is a canonical Clade III AAA+ protein and plays a regulatory role in the cellular processes of *B. anthracis*. No kinase activity was detected in the His6-PrkA, likely due to the lack of P-loop in its kinase domain (Supplementary figure-1A-C), as reported previously [11]. Furthermore, as shown in Figure-1A-D, purified His6-PrkA displayed negligible protease activity against substrates like LITAF and PA, even after incubation for 24 hours. Size-exclusion chromatography coupled with MALS (SEC-MALS) [29] revealed a monomeric form of purified His6 PrkA. Notably, an earlier preparation done in buffers without CHAPS and TCEP contained trace amounts of an amido-hydrolase contaminant (<0.1%) and had very weak protease activity. Similar issues may explain previously reported low-level protease activity in *Bacillus subtilis* PrkA (Supplementary Figure 5) [11]. Upon re-purification under reducing conditions, BAS PrkA had to detectable protease activity. These findings highlight the need for stringent purification to avoid misleading enzymatic activity and suggest that *Bacillus anthracis* PrkA may exert its regulatory effects through non-enzymatic mechanisms.

Functional analyses revealed that deletion of BAS *prkA* in *B. anthracis* Sterne 34F2 compromises growth under osmotic stress and impacts the transcription of key sporulation-related genes, resulting in defective spore coats (Figure-2-4). Although BAS *ΔprkA* progresses through the sporulation stages with the same morphological appearance as the wild-type strain, it exhibits defects in spore-coat formation resulting in heat sensitivity in the mature BAS *ΔprkA* spores.

Gene expression analysis revealed significant downregulation of *cotE* and *gerK*, which are critical for coat formation and germination [30, 31], respectively. Transcriptional analysis of several genes regulated by sporulation-related sigma factors, *sigF*, *sigG*, *sigE*, and *sigK,* confirms that PrkA exerts pleiotropic effects during sporulation. This is likely due to reduced expression of *sigK* or its downstream genes (*cotE*, *spoIVA*, and *spoIVB*), rather than through direct enzymatic activity. The downregulation of other essential sporulation genes like *spoIID, spoVR and gerK* in BAS *ΔprkA* mutant also suggest that PrkA has broad effect on sporulation pathway.

To elucidate the mechanistic basis of BAS PrkA’s role in *B. anthracis* life cycle, we mapped its interactome through co-affinity purification and mass spectrometry (Figure-5). The proteins identified as potential interacting partners interactors of PrkA are involved in septum formation, metabolic pathways, stress responses, and gene regulation. Notably, ProA, EzrA, and Ppx emerged as key interactors linked to the observed phenotypes. ProA, a central enzyme in proline biosynthesis, supports osmotic stress adaptation by maintaining osmotic balance [25, 26]. EzrA, a septation ring regulator, is essential for cell division and may have a novel role in spore maturation and viability [27]. Ppx, implicated in stress responses and sporulation efficiency [28], further connects PrkA to stress adaptation mechanisms. The absence of proteolytic activity in BAS PrkA suggests an alternative mechanism of action, potentially acting as a competitive inhibitor to other AAA+ proteins or as a molecular scaffold. Such proteins are known to regulate processes by binding substrates or cofactors without catalysis [32, 33] and this paradigm may extend to BAS PrkA. By interacting with key players like ProA, EzrA, and Ppx, PrkA could serve as a regulatory hub, modulating sporulation and stress response pathways in *B. anthracis*.

## Conclusion

In summary, this study suggests BAS PrkA potentially acting as a competitive inhibitor to other AAA+ proteins or molecular scaffold in *B. anthracis*, with essential roles in sporulation and stress adaptability. Our findings provide new insights into the molecular mechanisms of spore development and stress response in *B. anthracis*, highlighting the evolutionary versatility of enzymatically dead proteins in bacterial systems.

## Materials and methods

### Bacterial strains and growth conditions

*E. coli* strains DH5α and SCS110 were used for cloning, and BL21(DE3) was used for the expression of recombinant proteins (Table 1). Cultures were grown in Luria Bertani (LB) broth (Difco) supplemented with appropriate antibiotics at 37°C with constant shaking at 200 rpm. Ampicillin and kanamycin were used at final concentrations of 100 µg/mL and 25 µg/mL, respectively.

*B. anthracis* Sterne 34F2 strain (BAS WT) was used as the background strain for the construction of all other strains (Table 1). All *B. anthracis* strains were grown in LB broth (Difco) at 37°C with constant shaking at 200 rpm or in sporulation media at 30°C with shaking at 200 rpm. Kanamycin was used at final concentrations of 15 µg/mL.

### Cloning expression and protein purification

The coding sequence of BAS0518 (BAS *prkA*) was PCR amplified using gene-specific forward and reverse primers (Table-2) containing BamHI and XhoI restriction enzyme sites at the 5’ and 3’ ends. The PCR product was cloned into pPro-Ex-HTc vector using the BamHI and XhoI restriction enzymes (NEB) followed by ligation using T4 DNA ligase (NEB). Ligation was transformed into *E. coli* DH5α and clones were sequenced using Sanger sequencing (Psomagen). Recombinant plasmids were subsequently introduced into *E. coli* BL21(DE3) cells for protein expression. A starter culture of the recombinant *E. coli* strain, containing the desired plasmid, was grown overnight in 5 mL of LB broth supplemented with ampicillin (100 μg/mL). This primary culture was then used to inoculate a 1-L secondary culture in LB broth with ampicillin (100 μg/mL), which was incubated at 37°C with shaking at 200 rpm. The culture was grown until the OD_600_ reached 0.6–0.8. At this point, protein expression was induced by adding IPTG to a final concentration of 1 mM, and the culture was incubated for an additional 3 hours at 37°C. Cells were then harvested by centrifugation at 8000 rpm for 5 minutes, and the resulting pellet was washed with 1X PBS.

The cell pellet was resuspended in sonication buffer [50 mM Tris-HCl (pH 8.5), 5 mM β-mercaptoethanol, 1 mM phenylmethylsulfonyl fluoride (PMSF), 1X protease inhibitor cocktail (Roche Applied Science, USA), and 300 mM NaCl] and subjected to sonication (9 cycles at 20 % amplitude, 10 seconds on, 30 seconds off). The lysate was then centrifuged at 13,000 rpm for 1 hour. The supernatant containing the recombinant protein was purified by immobilized metal affinity chromatography (IMAC) and eluted with 200 mM imidazole. The purified proteins were pooled, dialyzed for downstream assays, and stored at -80°C.

For purification of GST-tagged catalytic domain of PrkC (PrkC^cat^-GST), the recombinant protein was purified using the method described earlier [14].

### Generation of single gene knockout strains in *B. anthracis*

The BAS *prkA* knockout strain in *B. anthracis* was generated using the method described by Pomerantsev et al. (2009). Briefly, two sequential single-crossover events were employed to insert *loxP* sites bracketing the gene to be targeted, andexpression of Cre recombinase then excised the gene, leaving a single loxP site [14, 20] . Specifically, the upstream (left) and downstream (right) regions of the BAS *prkA* gene were PCR amplified with primers listed in Table-2. Subsequently the PCR amplicons was cloned (separately) into the SpeI and XhoI sites of the pSC vector (described below). The resulting plasmids, 0518-left-pSC and 0518-right-pSC, were passed through the *dam*^-^/*dcm*^-^ *E. coli* strain SCS110 to ensure the plasmids were unmethylated. These plasmids were then sequentially electroporated into *B. anthracis* Sterne 34F2, and transformants were selected on LB agar plates supplemented with erythromycin (10 μg/mL).

Subsequent transformation with pCrePAS2 (described below) induced Cre recombinase expression, then 0518-right-pSC and finally a repeat of pCrePAS2 resulting in the excision of the DNA region between the two *loxP* sites. Approximately 20 colonies were screened for sensitivity to both spectinomycin and erythromycin. Colonies that were resistant to neither antibiotic were further confirmed by PCR using primers flanking primer (0518-LseqF and 0518-LseqR, Table 2).

**Table 2:**
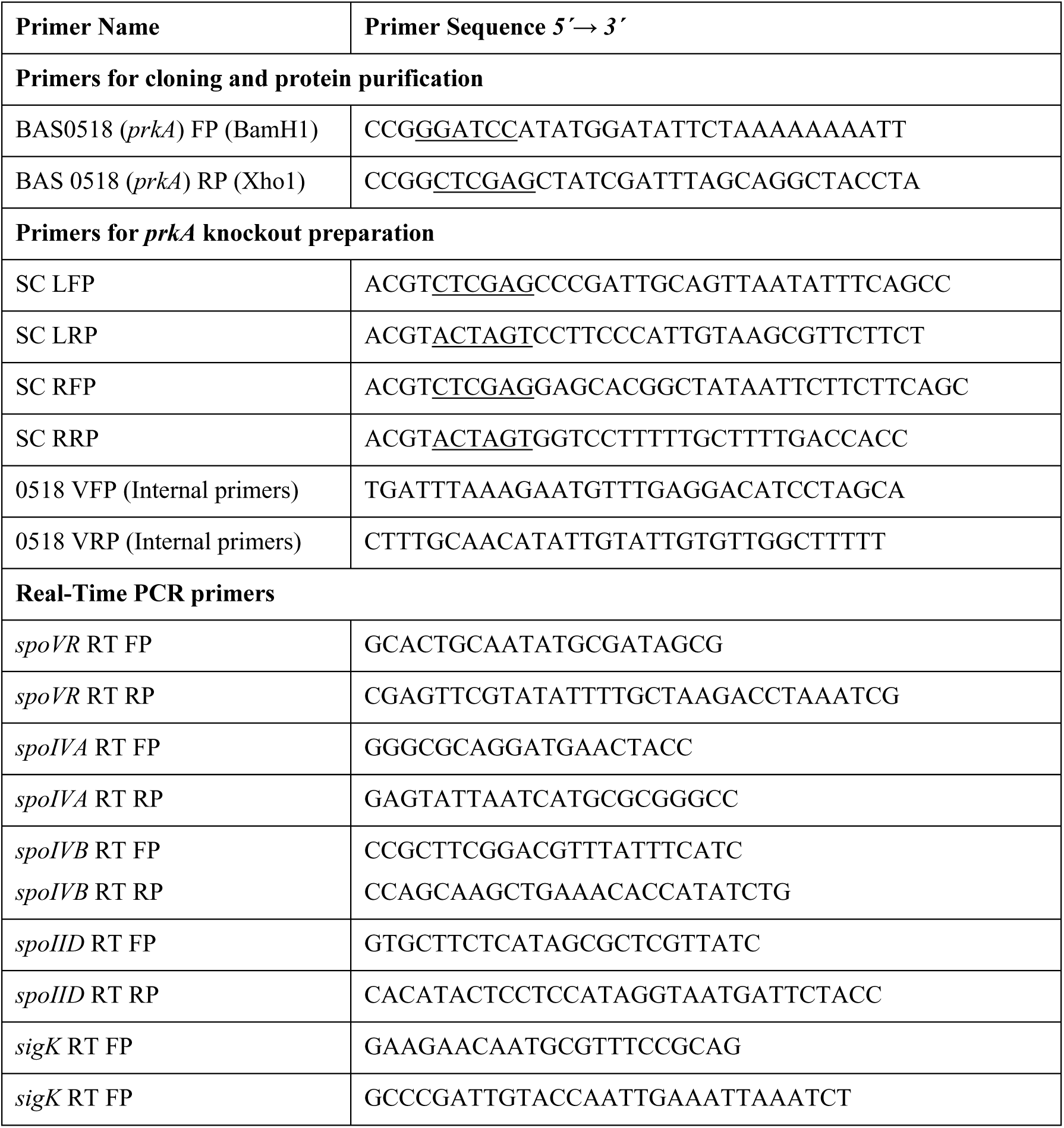
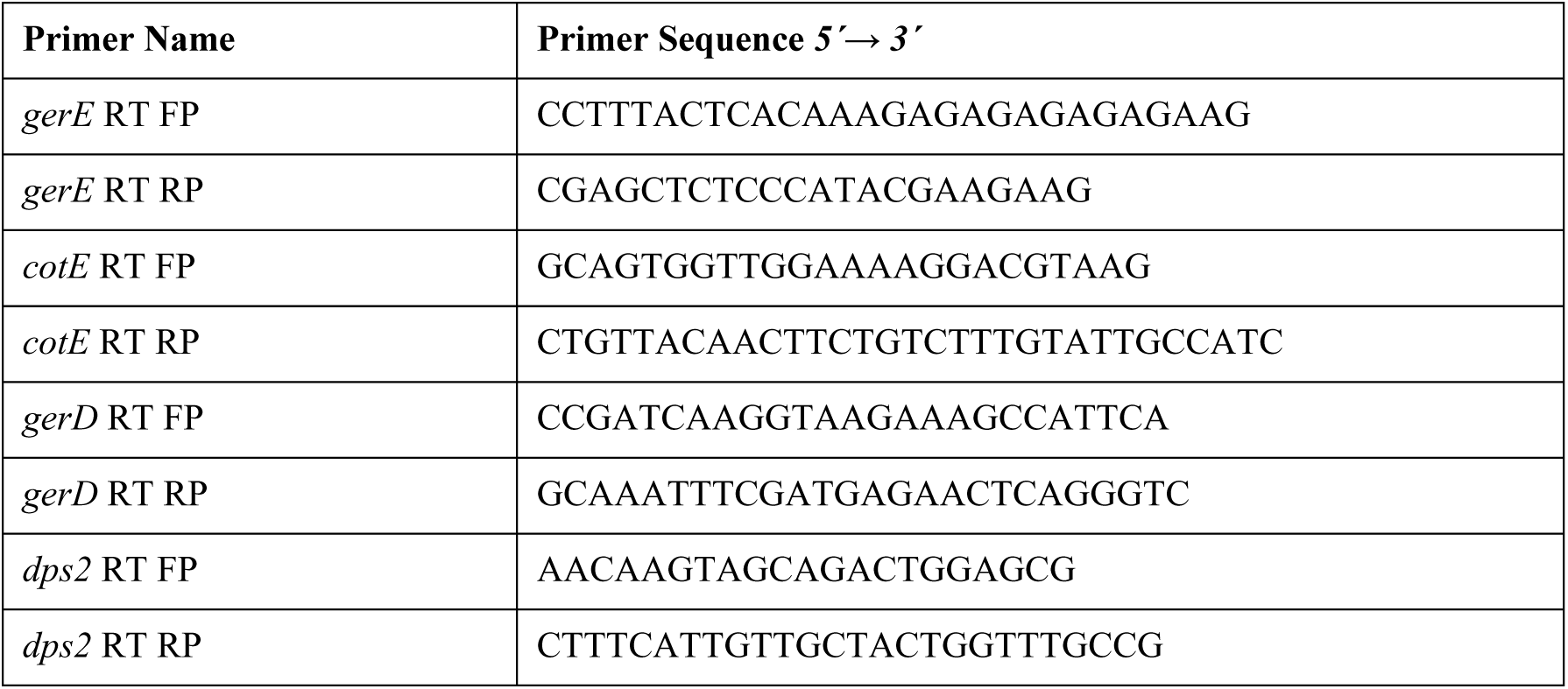
List of primers used for cloning, gene deletion and qPCR analysis:

### Generation of Complement strain of BAS *prkA*

The coding sequence of BAS *prkA* was synthesized as gBlock (IDT) with homologous ends to the pAMY1 vector digested with HindIII/SalI. Using NEB HiFi, the gBlock was cloned into the digested pAMY1 to generate pAMY-*prkA* (Table 1). The complemented strain (BAS *ΔprkA*::pAMY*-prkA*) was generated by electroporating the BAS *ΔprkA* strain with pAMY-*prkA* plasmid.

### Growth kinetics

The primary culture of BAS WT and BAS BAS *ΔprkA* strains were grown in LB broth at 37°C and 200 rpm overnight. The overnight culture was added as the inoculum to begin secondary cultures in triplicates at starting OD_600_ of 0.05 in similar growth conditions. Absorbance at 600 nm was recorded throughout the experiment after every hour until the cells reached the stationary phase.

### PrkA phosphorylation assay

Autophosphorylation activity of His6-PrkA was assessed using an anti-phosphothreonine (anti-pThr) antibody, following a previously described method [34]. Briefly, 0.5 µg/well of His6-PrkA was incubated with or without 1 mM ATP in a reaction buffer containing 20 mM HEPES (pH 7.4), 5 mM MgCl_2_, 5 mM MnCl_2_, 1 mM DTT for 30 minutes at 25°C.

As a positive control for kinase activity, we used the GST-tagged catalytic domain of PrkC (PrkCcat-GST), along with purified elongation factor Tu (Ef-Tu), myelin basic protein (MBP), and GST as substrates [34]. For each indicated lane, 0.5 µg/well of PrkCcat-GST was combined with 5 µg/well of Ef-Tu, 10 µg/well of MBP, or 5 µg/well of GST. Each reaction was incubated under the same conditions described above. To test whether PrkA could serve as a substrate for another kinase, His₆-PrkA was also incubated with PrkCcat-GST under identical assay conditions.

Reactions were resolved on a 4–20% SDS-PAGE gel, and proteins were transferred onto a nitrocellulose membrane using wet transfer at 4 °C for 2 hours at 120 V. Membranes were probed with rabbit anti-pThr (Thermo Fischer) and rat anti-PrkA (see below) primary antibodies, followed by Rabbit IgG (H&L) Antibody DyLight™ 700 Conjugated (Rockland) and Rat IgG (H&L) Antibody DyLight™ 800 Conjugated (Rockland) was used as the secondary antibody for immunoblotting. Imaging was performed using a LI-COR Odyssey CLx system, and band intensities were quantified using Image Studio software (LI-COR).

### Protease Assay

Protease activity of His6 PrkA was assessed using two methods: 1) Fluorescently labelled substrate (α-casein) and 2) Visual analysis on SDS-PAGE.

For the first method, reactions were set up with 0.1 μg/μL of His6 PrkA and 0.1 μg/μL of substrates (α-casein, LITAF, and PA) in 1X digestion buffer (40 mM Tris/HCl, pH 7.5, 1 mM MgCl₂, and 1 mM ATP) and incubated at 37°C for 16 or 24 hours. Controls were prepared with enzyme or substrate alone. Reactions were terminated by adding 5X SDS-sample buffer and analysed by 12.5 % SDS-PAGE.

In the second method, protease activity was measured using the EnzChek™ Protease Assay Kit. BODIPY FL-Casein served as the substrate, and reactions were set up in Corning^TM^96-well solid black flat-bottom plates. Each reaction (100 µL) contained 1 µg of His6 PrkA in 1X digestion buffer. The protease activity was measured by adding 100 µL of a 10 µg/mL working solution of BODIPY casein. The plate was incubated at room temperature for 2-24 hours, and fluorescence was measured at 485 nm excitation/535 nm emission.

### Size-exclusion chromatography coupled with multi-angle light scattering (SEC-MALS)

To assess the native of His6 PrkA, protein was injected onto HighLoad 16/60 Superdex 200 column AKTA Pure HPLC system pre-equilibrated in buffer containing 200 mM Tris-HCl pH 8.0, 300 mM NaCl, 5mM CHAPS, 1 mM TCEP and 10 % glycerol. The HPLC system was connected to a DAWN HELEOS II detector featuring a quasi-elastic light scattering module, along with an Optilab T-rEX refractometer (Wyatt Technology). Data was analyzed using the ASTRA 8.1.2.1 software (Wyatt Technology Europe) and the different protein fractions were c

### Growth under stress conditions

To assess the growth of BAS WT and BAS *ΔprkA* under various stress conditions, including osmotic stress (0, 0.05, 0.1, 0.5, and 1 M NaCl), non-ionic osmotic stress, and oxidative stress (0, 0.1 mM, 1 mM and 10 mM H₂O₂), an endpoint assay (with 16-hour incubation) was performed. Both strains were grown overnight at 37°C with shaking at 200 rpm. The following day, stress conditions (osmotic stress with NaCl) were applied, and growth was monitored overnight in a 96-well plate.

To determine the number of viable cells, resazurin dye (R&D systems) was added at a 1:10 ratio, and fluorescence was measured at an excitation/emission of 540/590 nm (Victor3).

### *B. anthracis* spore preparation and spores’ quantification

The BAS WT-pAMY, BAS *ΔprkA*-pAMY, and BAS *ΔprkA*::pAMY-*prkA* strains were grown overnight in LB broth at 37°C with shaking at 200 rpm. The overnight cultures were spread onto LB-agar plates containing 15 µg/mL kanamycin and 250 µM IPTG. Plates were incubated at 37°C for 24 hours and then shifted to 30°C for 9 days to induce sporulation. Spore formation was monitored using DIC microscopy. Spores were harvested by resuspending them in 10 mL of deionized water, followed by washing three times at 12,000 x g for 15 minutes with sterile deionized water. The spores were then re-suspended in 5 mL of sterile deionized water. Half of the spore preparation was heat-treated at 75°C for 30 minutes, while the remaining half was kept at room temperature in sterile deionized water.

Spore viability was assessed by plating 100 µL of 10-fold serial dilutions of both heated and unheated spores on LB-agar plates containing 15 µg/mL kanamycin and 250 µM IPTG.

### PrkA antibody generation and temporal expression using immunoblotting

Anti PrkA antibodies were produced in female Fischer (CDF) rats (Charles River) by priming with 50 µg of purified His6 PrkA (antigen), followed by two boosts with 25 µg of antigen at two weeks and five weeks post-priming. All immunizations were by subcutaneous route and antigen was pre-mixed with an equal volume of Alhydrogel (Invivogen). Rats were bled 2 months post-priming and tested for anti PrkA antibody titters by standard ELISA. All animal immunizations were done in accordance with protocols approved by the NIAID ACUC (animal protocol LPD8E).

For the temporal expression of PrkA, 5 mL cultures of bacteria in vegetative and sporulating stages were collected and lysed in 1 mL lysis buffer [6 M Urea, 50 mM NH_4_CO_3_, 1 mM DTT] by performing 7 cycles of bead beating with 0.1 mm Zirconia Beads (BioSpec) using Precellys evolution homogenizer (Bertie technologies) at 9000 rpm, with 20 seconds ON and 1 minute OFF. The lysate was centrifuged at 13000 rpm for 30 minutes, and total protein concentration was estimated using the BCA method (Pierce, Invitrogen). Since BAS PrkA expression was tested in different stages of growth and sporulation, lysate volumes were normalized to the total protein concentration, and 20 µg of total lysate from each condition was resolved on a 4-20 % SDS-PAGE gel (Invitrogen). The proteins were transferred onto a nitrocellulose membrane using the iBlot2 (Invitrogen). The blot was blocked and probed using rat polyclonal anti-PrkA at a dilution of 1:5000 in LI-COR blocking buffer. Rat IgG (H&L) Antibody DyLight™ 800 Conjugated (Rockland) was used as the secondary antibody for immunoblotting, and blots were scanned on the LI-COR Odyssey CLx.

### Fluorescent microscopy for spores and sporulation stages

Sporulation stages in BAS WT and BAS *ΔprkA* were monitored using the cell-permeable dye hexidium iodide at 100 µg/mL. Briefly, bacterial cells were synchronized by streaking single colonies of each strain onto LB-agar plates and incubating at 37°C for 16 hours. Single colonies from each strain were then inoculated into 5 mL sporulation media (8 g LB broth/L supplemented with 85.5 mM NaCl, 0.025 mM ZnSO₄, 0.6 mM CaCl₂, 0.3 mM MnSO₄, 0.8 mM MgSO₄, and 0.02 mM CuSO₄, pH 5.0 -5.5) and incubated at 30°C with shaking at 220 rpm to induce sporulation.

At different time points between 13-16 hours post-inoculation, 200 µL of each culture was collected, washed with 1X PBS, and incubated with 100 µg/mL hexidium iodide for 20 minutes at room temperature. The cells were then fixed with 4 % paraformaldehyde in 1X PBS overnight at 4°C. Fixed cells at different sporulation stages were imaged using a Leica SP8 microscope (690/730).

For the hexidium iodide (HI) diffusion assay in spores, WT-pAMY, BAS *ΔprkA*-pAMY, and BAS *ΔprkA*::pAMY-*prkA* strains were used. Spores were prepared as described above, stained with 100 µg/mL hexidium iodide for 20 minutes, and fixed with 4 % paraformaldehyde in 1X PBS overnight at 4°C. HI-stained spores were imaged using the Leica SP8 microscope (690/730). Spores exhibiting surface staining were considered viable, while those with diffuse staining were classified as non-viable. The number of surface-stained and diffused-stained spores was manually counted.

### RNA extraction and quantitative real-time PCR

BAS WT and BAS *ΔprkA* strains were grown in triplicates to the early stationary phase, and RNA was extracted using a modified hot lysis method as previously described [35–37]. Briefly, cells harvested at the desired growth stage were resuspended in 500 µL TRIzol® (Invitrogen). To this, 400 µL of hot lysis buffer (50 mM Tris, pH 8.0, 1 % SDS, 1 mM EDTA) and 400 µL of DEPC-treated Zirconia beads were added. The cell suspension was incubated at 65°C for 15 minutes with intermittent vortexing every 5 minutes. After cooling on ice, 100 µL of chloroform per mL of TRIzol was added, and the suspension was centrifuged at 9,500 × g for 15 minutes at 4°C to separate the RNA in the aqueous phase. RNA was precipitated by adding 0.5 M LiCl₂ and 3X ice-cold isopropanol, followed by incubation at -80°C for 2 hours. The RNA was pelleted by centrifugation at 16,000 × g for 20 minutes at 4°C, washed with 70 % ethanol (Merck), and resuspended in nuclease-free water after air-drying. DNA contamination was removed by treating the sample with DNase (Ambion) as per the manufacturer’s instructions. The purified RNA was then used for cDNA synthesis using a first-strand cDNA synthesis kit (Thermo Fisher).

To analyse the expression of sporulation genes at different growth phases (log, late exponential/early stationary, and late stationary), cDNA was synthesized from the RNA samples and analysed using gene-specific primers and SYBR Green master mix (Roche). Reactions (10 µL each) were run in triplicates with no-template controls in a Quant Studio 7 (Thermo Fisher). The housekeeping gene *rpoB* (encoding DNA-directed RNA polymerase subunit beta) [38] was used for normalization. The primers used in this study were designed to amplify PCR products of 100–150 bp.

### Co-affinity purification and mass spectrometry

*B. anthracis* strains were cultured in 200 mL of LB broth. The strains used included BAS WT-pAMY and BAS WT-pAMY-*prkA*-TEV-His6. Once cultures reached an OD_600_ of ∼ 0.2, 50 µM IPTG was added, and growth continued until the cultures reached an OD_600_ of ∼ 2. Cells were harvested by centrifugation at 6,200 × g for 10 minutes at 4°C, and the wet weights were recorded. Pellets were snap-frozen on dry ice and stored overnight. The following day, the cells were thawed in an ice-water bath and resuspended in 50 mL of binding buffer (5 mM imidazole, 0.5 M NaCl, 5 mM β-mercaptoethanol, 20 mM Tris pH 7.2) supplemented with an EDTA-free protease inhibitor cocktail. Cells were lysed by two rounds of French press, and the soluble fraction was collected by centrifugation at 10,000 × g for 10 minutes at 4°C.

For co-purification, the soluble material was clarified by an additional centrifugation at 18,000 rpm for 10 minutes at 4°C, followed by a 20-minute incubation at 37°C. In the meantime, 1 mL of Ni-NTA resin was equilibrated with 50 mL of binding buffer for 20 minutes. The equilibrated Ni-NTA resin was then added to the samples, and the mixture was incubated at 4°C for 2 hours to allow His6-tagged proteins to bind to the resin. Following incubation, the resin was transferred to a column and washed at room temperature by gravity flow using the following steps-three rounds of 5 mL binding buffer, two rounds of 5 mL Wash Buffer 1 (40 mM imidazole pH 7.9, 1.0 M NaCl, 20 mM Tris pH 8.0, 5 mM β-mercaptoethanol), one round of 10 mL Wash Buffer 1, one wash with 1.5 mL High Salt Wash Buffer (40 mM imidazole pH 7.9, 1.5 M NaCl, 20 mM Tris pH 8.0, 5 mM β-mercaptoethanol, 5 mM CHAPS), and three rounds of 2.5 mL Wash Buffer 1. His6-tagged PrkA and its interacting proteins were eluted in three 500 µL aliquots of elution buffer (20 mM Tris pH 7.5, 300 mM imidazole pH 7.9, 500 mM NaCl, 5 mM β-mercaptoethanol). The eluted samples were analysed by SDS-PAGE and sent to the NIAID Research Technology Branch (RTB) core facility for protein identification via mass spectrometry.

### Pathway enrichment analysis

ShinyGO 0.81 [24] was used to do pathway enrichment analysis of the BAS0518 interacting proteins. UniProt was used as the pathway database with species name set to *B. anthracis* str. ’Ames Ancestor’ (Accession number NC_007530). An FDR cutoff of 0.05 was used to identify significantly enriched pathways.

### Statistical Analysis

Graphs and statistical analysis were performed using GraphPad Prism. P-value is calculated using two-tailed Student’s T test.

## FUNDING AND ACKNOWLDGEMENT

This research was supported by the intramural research program of the National Institute of Allergy and Infectious Diseases of National Institutes of Health. Nitika Sangwan was supported by CSIR-NET fellowship from Government of India and Graduate Partnership Program from NIH. We would like to thank RTB staff for their support with mass-spectroscopy and microscopy.

## AUTHOR’S CONTRIBUTION

**Nitika Sangwan:** Performed experiments (cloning, gene deletion, protein purifications, microscopy assay, AlphaFold), validation (biochemical assays, immuno-blotting), formal analysis, investigation, visualization, manuscript writing. **Ankur Bothra:** Conceptualization, supervision, methodology (cloning, gene deletion, protein purifications, microscopy assay, AlphaFold), validation (biochemical assays, immunoblotting), formal analysis, investigation, visualization, and manuscript writing. **Aakriti Gangwal**: Input and discussion for experiments. **Andrei Pomerantsev**: Gene deletion, manuscript editing. **Rasem Fattah:** Protein purifications. **Mahtab Moayeri**: Antibody generation, manuscript editing. **Sundar Ganesan**: Microscopy assay. **Qian Ma:** Animal handling and antibody generation. **Chetkar Chandra Keshavam:** Manuscript editing. **Renu Baweja**: Discussion and Manuscript editing. **Uma Dhawan**: Conceptualization, supervision, and manuscript editing. **Stephen H. Leppla**: Manuscript editing, resources, supervision and funding acquisition. **Yogendra Singh**: Conceptualization, supervision, manuscript editing, resources, and funding acquisition.

